## Supplementary figures and images for "Dominant spinal muscular atrophy linked mutations in the cargo binding domain of BICD2 result in altered interactomes and dynein hyperactivity"

### Supplemental figure1

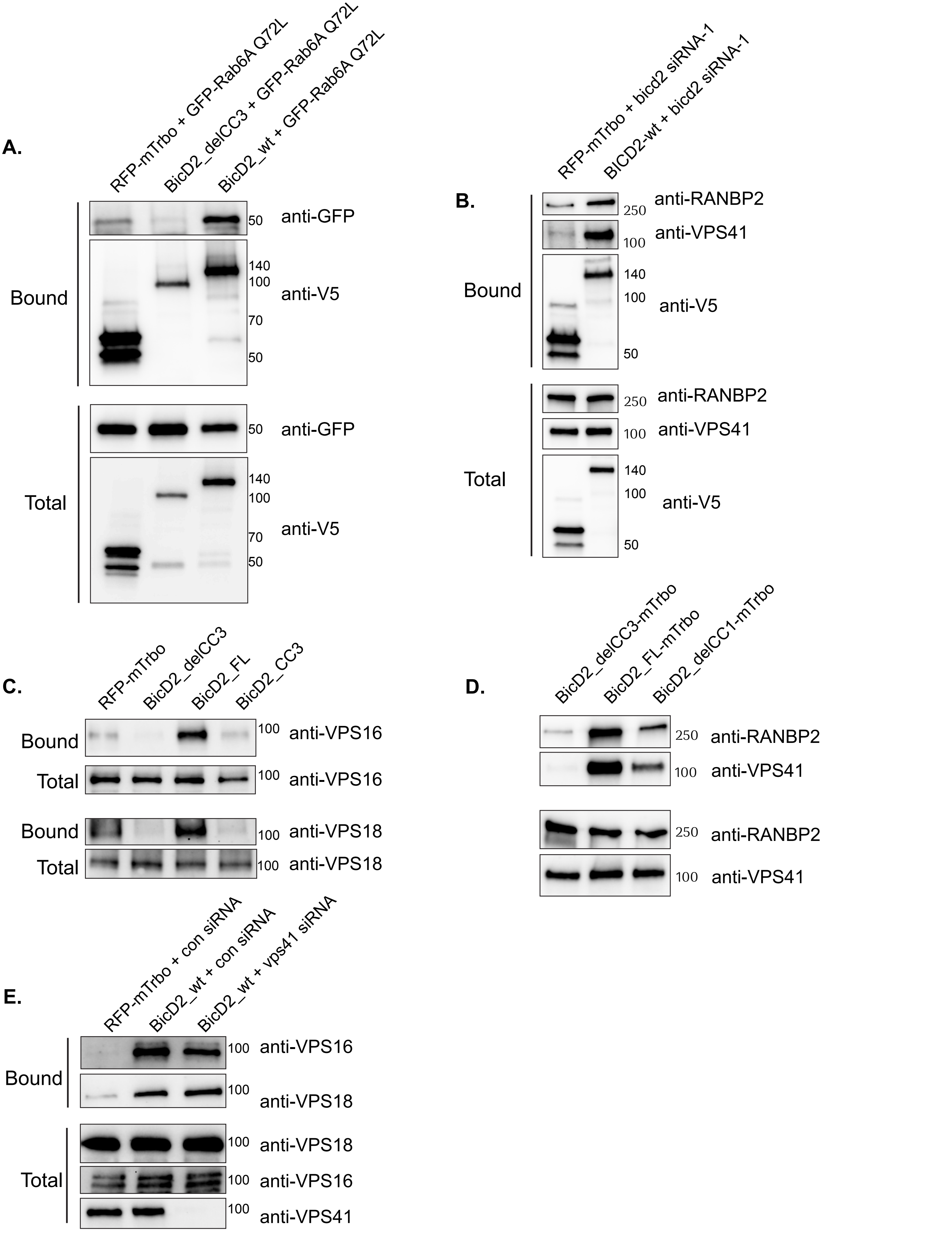


Supplemental Figure 1

### Supplemental figure2

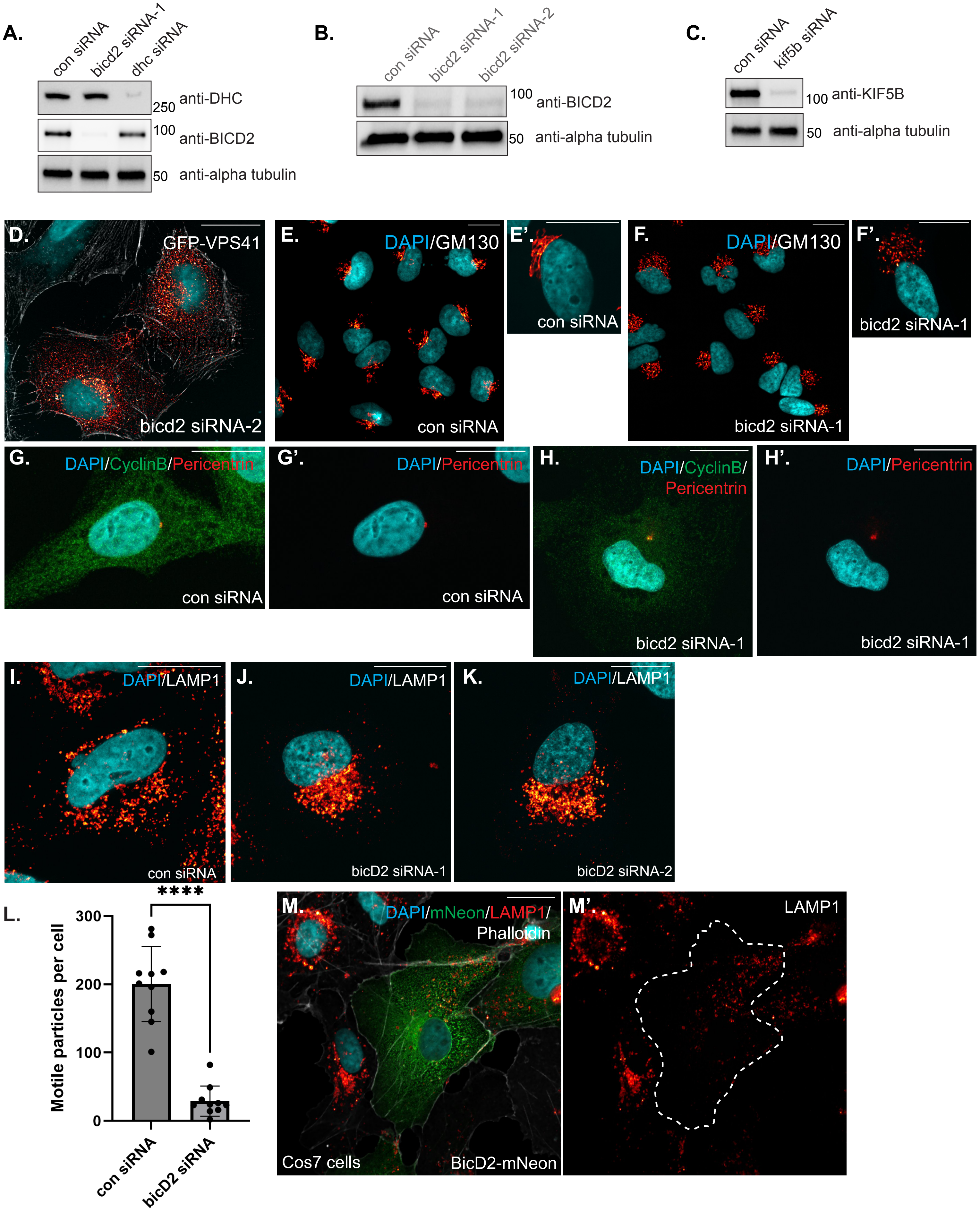

### Supplemental figure3

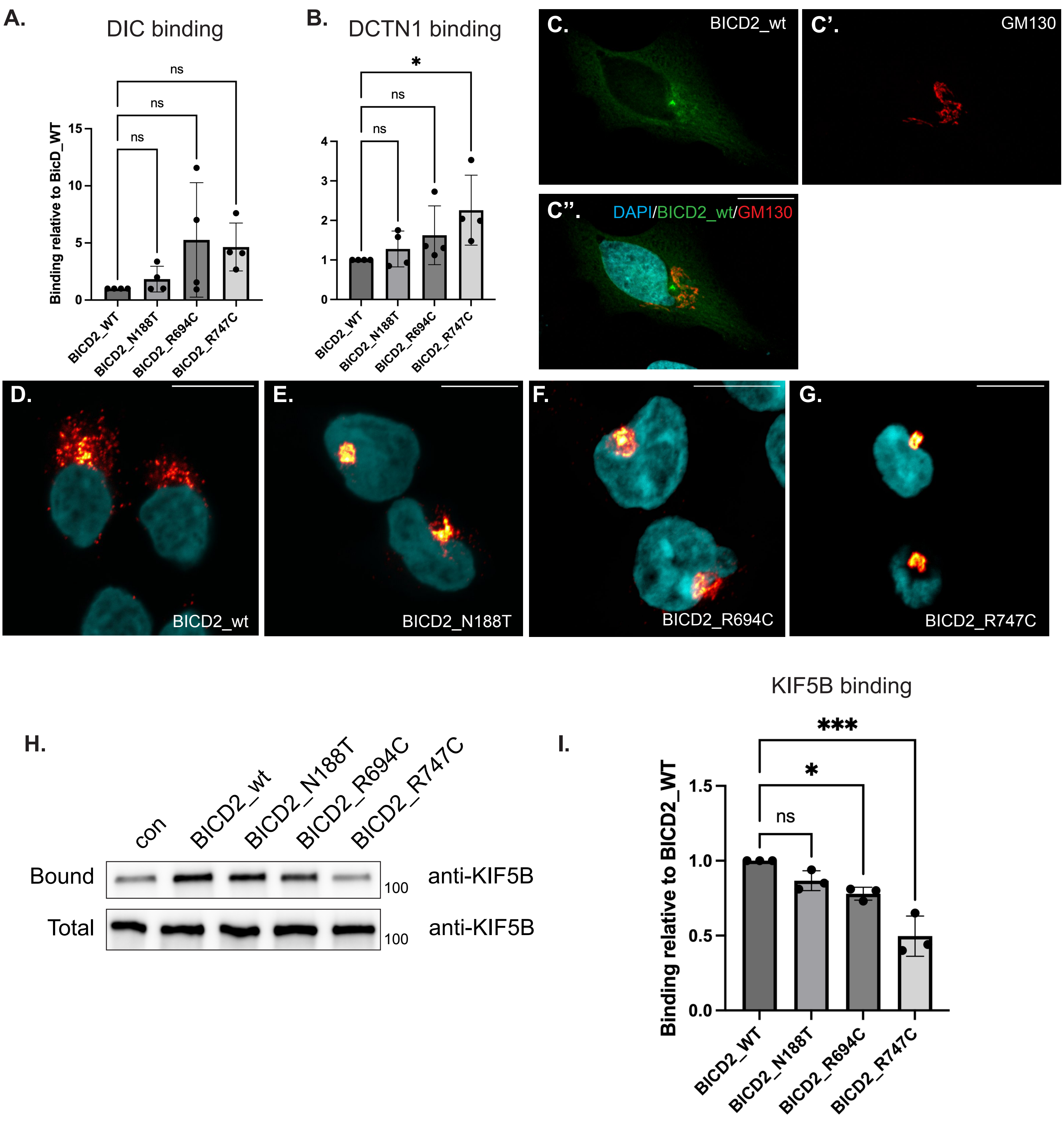


Supplemental Figure 3

### Supplemental figure4

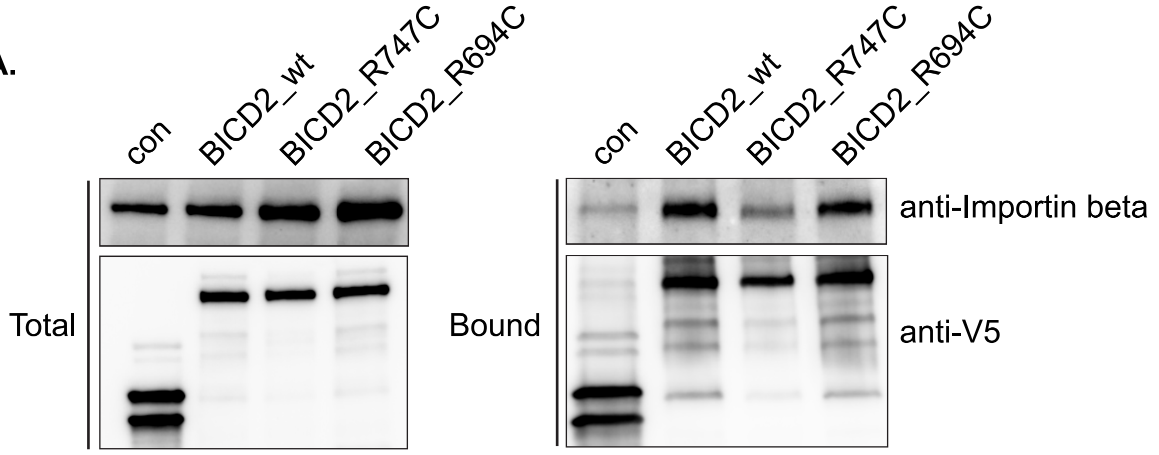

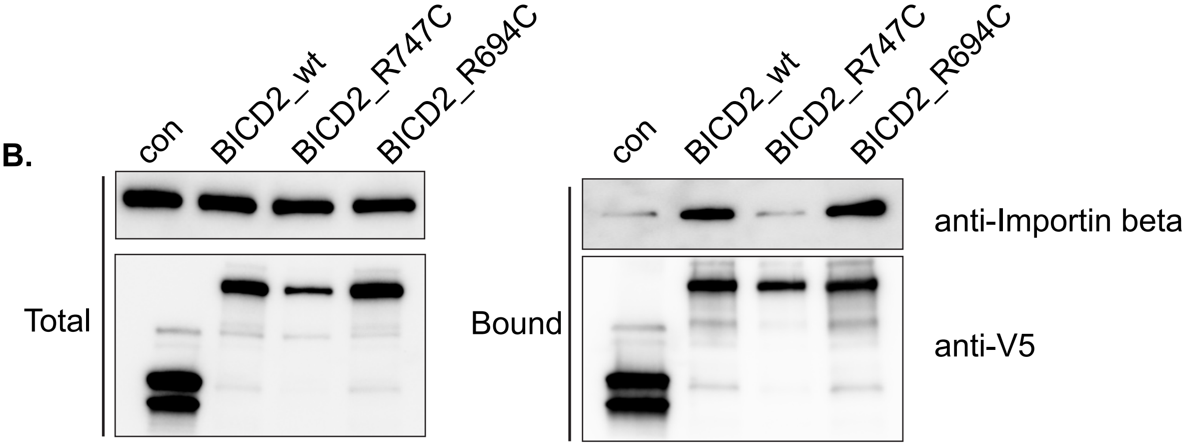

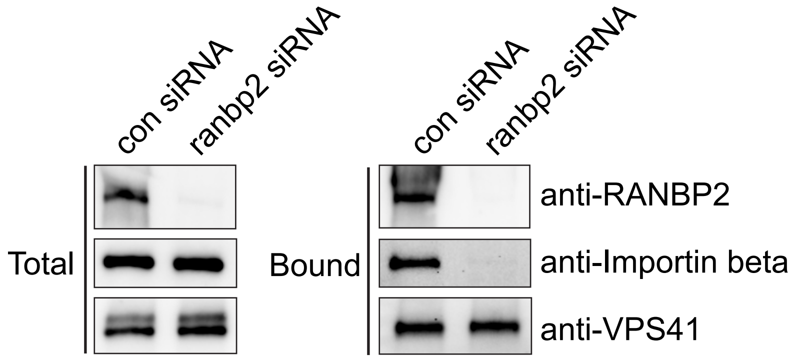

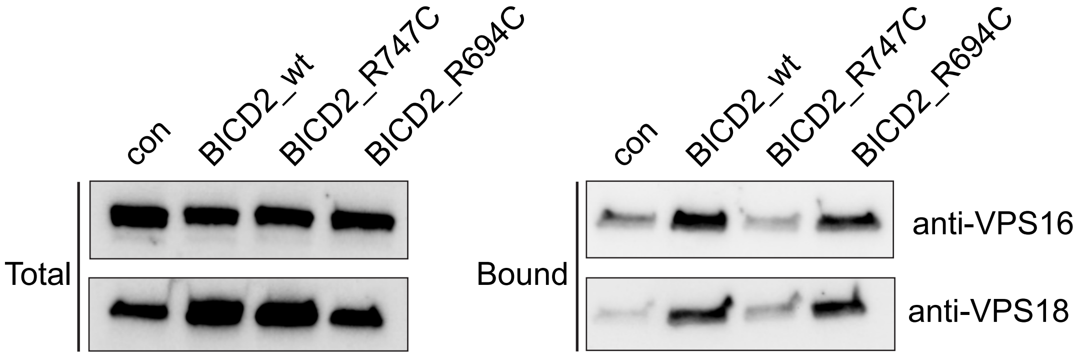

### Supplemental figure5

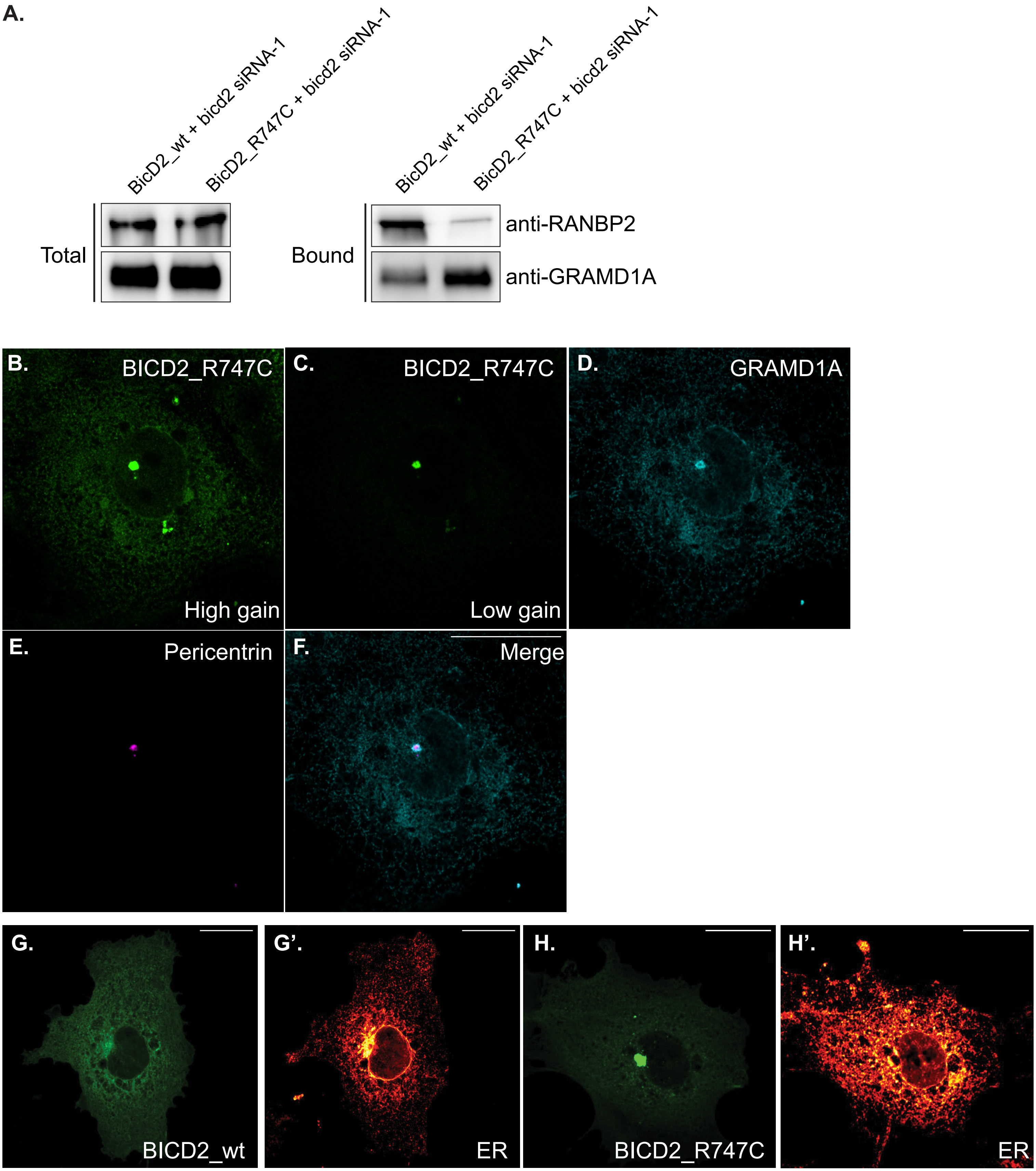
